# Tetrapods on the EDGE: Overcoming data limitations to identify phylogenetic conservation priorities

**DOI:** 10.1101/232991

**Authors:** Rikki Gumbs, Claudia L. Gray, Oliver R. Wearn, Nisha R. Owen

## Abstract

The scale of the ongoing biodiversity crisis requires both effective conservation prioritisation and urgent action. The EDGE metric, which prioritises species based on their Evolutionary Distinctiveness (ED) and Global Endangerment (GE), relies on adequate phylogenetic and extinction risk data to generate meaningful priorities for conservation. However, comprehensive phylogenetic analyses of large clades are extremely rare and, even when available, become quickly out-of-date due to the rapid rate of species descriptions and taxonomic revisions. Thus, it is important that conservationists can use the available data to incorporate evolutionary history into conservation prioritisation. We compared published and new methods to impute ED for species missing from a phylogeny whilst simultaneously correcting the ED scores of their close taxonomic relatives. We found that following artificial removal of species from a phylogeny, the new method provided the closest estimates of their “true” score, differing from the true ED score by an average of less than 1%, compared to the 31% and 38% difference of the previous imputation methods. Previous methods also substantially under- and over-estimated scores as more species were artificially removed from a phylogeny. We therefore used the new method to estimate ED scores for all tetrapods. From these scores we updated EDGE prioritisation rankings for all tetrapod species with IUCN Red List assessments, including the first EDGE prioritisation for reptiles. Further, we identified criteria to identify robust priority species in an effort to further inform conservation action whilst limiting uncertainty and anticipating future phylogenetic advances.

## Introduction

We are currently in a period of unprecedented human-mediated biodiversity loss, often termed the ‘sixth mass extinction’ [1]. To achieve global commitments to halt the loss of biodiversity [2], the resources available for conservation must be targeted effectively. Several species-level conservation prioritisation schemes [3] have been developed, focussing on ‘charismatic’ species [4,5], threat levels [6], important ecosystem service providers [7], or some combination of these [8–11].

However, very few—if any—of these approaches explicitly focus on preserving unique evolutionary history, or Phylogenetic Diversity (PD) [12–14]. Species with few extant close relatives represent a disproportionate amount of the total PD of their clade [15]. Where these species are threatened with extinction, they often represent a significant amount of important trait diversity that could soon be lost [16,17]. Therefore, current conservation prioritisation approaches that do not take PD into consideration may fail to prevent the loss of large amounts of both phylogenetic and trait diversity [13,17–19]. To date, several metrics have been proposed to integrate PD into the prioritisation of species and regions [12,15,20–23].

A number of these metrics measure the contribution of individual species to the total PD of a clade [24–29], and the Evolutionary Distinctiveness (ED) metric has received the most widespread use [12,14,15,30–35]. Whereas PD is the sum of all branch lengths of a phylogeny, ED is the proportion of the total PD assigned to an individual species, with the length of each branch of the phylogeny divided equally amongst all species to which it is ancestral (see original formulation [12] for detailed description). This partitioning of PD amongst species facilitates prioritisation at the species, rather than clade, level.

In conjunction with PD prioritisation, conservation actions must also be timely. Many species are at imminent risk of extinction, and those that are under greatest threat are widely considered to be the highest priority for immediate action. The EDGE metric, which combines the ED of a species with its extinction risk—or ‘Global Endangerment’ (GE) [12], has been implemented by the EDGE of Existence programme at the Zoological Society of London to prioritise species in a number of taxonomic groups (mammals [12,19], amphibians [30], birds [15], and corals [36]). The EDGE of Existence programme is the only global conservation initiative to focus on threatened species representing a significant amount of unique evolutionary history, raising awareness of these often poorly known species, and actively supporting conservation activities [37]. Research has shown the EDGE metric has the potential to not only preserve more PD than expected [17], but also preserve more trait diversity than expected if conserving threatened species without considering PD [16,17].

However, meaningful and credible prioritisation for conservation depends on the quality of data available. Metrics such as ED ideally require species-level phylogenies to calculate the individual contribution of each species to the total PD of a clade [12,32], yet no phylogeny exists that contains all known species of any tetrapod class. There are little, if any, genetic data available for many poorly-known species, precluding their inclusion in most phylogenetic analyses. In addition, given the high rate of tetrapod species descriptions, species-level phylogenies quickly become out of date; for example, almost 400 species were missing from the mammalian supertree [38] less than four years following publication [19]. Even though the exact phylogenetic position of these “missing species” is not known, in most cases they can be assigned to genus or family [36,39,40]. This provides an opportunity for their ED scores to be estimated based on the ED scores of congeneric or confamilial species.

The phylogenies currently available also have notable limitations. Of the tetrapods (amphibians, birds, mammals and reptiles), amphibians and testudines suffer from particularly poor phylogenetic coverage [41–44], reflecting the relatively low research investment in these taxa compared to birds and mammals [45–47]. For example, at the time of writing the largest published amphibian genetic phylogeny [41] omits more than 3,600 species (50% of known species). Recent species-level phylogenies published for birds [48], mammals [38,49,50], and squamates [35] represent advances in phylogenetic coverage for these groups, but many species are still missing (~1,000 birds, ~500 mammals, ~200 squamates respectively). To overcome paucity of genetic data, many phylogenies are now constructed using taxonomic information and constraints to infer phylogenetic relationships for species lacking available genetic data [40,48,51]. Such phylogenies are inherently uncertain and therefore produce a large distribution of equiprobable phylogenetic trees, rather than a single consensus phylogeny, in order to capture the uncertainty around taxonomically-inferred relationships [40,51]. This reliance on taxonomic data means existing phylogenies are susceptible to significant changes with more comprehensive genetic sampling [40,50,51].

The uncertainty in available phylogenies must be accounted for and acknowledged when developing conservation priorities. Given the imminent biodiversity crisis [1], it is impractical and undesirable for conservationists to wait for completely inclusive phylogenies to be published before implementing PD-based conservation efforts [19]. We therefore required a reliable method for incorporating all known species when using incomplete or out-of-date phylogenies.

Two statistical imputation methods have previously been employed to calculate ED for species missing from phylogenies [12,19,36], though the relative performance of these methods has not yet been examined. Here, we compare the accuracy of both of the existing imputation methods with that of a third, novel method. We show empirically that the ED, and also EDGE rank, of missing species can be accurately predicted from other species in the phylogeny using our new method. This produces a robust set of priority species, and deals effectively with the uncertainty inherent in phylogenetic data. Finally, we use the statistical imputation of ED scores to produce updated EDGE priority lists for all tetrapods, including the first EDGE list for reptiles.

## Materials and methods

### Imputation of Evolutionary Distinctiveness for missing species

Isaac et al.’s [12] method of imputing ED scores for missing species (hereafter the ‘original’ imputation method) reduced the ED scores of species in the phylogeny in proportion to the number of missing congeners or confamiliars, and assigned the difference in ED scores between the original and corrected values for each species present in the phylogeny evenly across the missing species. This reduction of ED scores for species with missing congeners or confamiliars replicates an inherent characteristic of ED calculations: the amount of ED received from an internal branch is inversely proportional to the number of descendant species [12]. However, as this method shares existing ED from species present in the phylogeny amongst the missing relatives without adding new ED to the phylogeny, it fails to replicate a second inherent characteristic of ED calculation: the addition of species with non-zero terminal branch lengths will invariably increase the total ED (= PD) of the clade, which would be expected with the inclusion of additional, missing, species [12].

Conversely, Curnick et al. [36] followed Swenson [39] by calculating the mean ED values for genera and families in the phylogeny and assigning this score to missing congeners and family members (hereafter the ‘simple’ imputation method). This imputation provides the opposite result of the ‘original’ imputation: though it does increase the PD of the clade with the inclusion of missing species, the simple method fails to reduce the ED scores of the closely-related species present in the phylogeny from which the scores are imputed.

We employed a novel method to estimate ED for missing species that both corrects the ED score of species present in the phylogeny and increases the PD of the clade. First, we calculated ED scores for all missing species using the ‘simple’ imputation method. Second, for genera with missing species and families with missing genera, we calculated the total ED for the associated species present in the phylogeny and divided this equally amongst all congeners or congeners, including those missing from the phylogeny. Each species now had two ED scores. For species in families with no species missing from the phylogeny, the two ED scores were identical. We then calculated the mean of the two values to derive an ED score that both increases the total PD of the clade upon inclusion of missing species and corrects the ED of species with missing relatives.

To assess the performance of the three imputation methods we took available dated consensus phylogenies for three tetrapod clades; amphibians [41], mammals [49], and squamates [52]. To simulate the imputation of ED scores from phylogenies that accurately represent all higher-level relationships but do not contain all species within a genus or genera within a family (e.g. [41,52,53]), we followed four steps for each phylogeny: 1) a ‘reference’ ED score for all species was calculated from the unaltered phylogeny; 2) a random number of species were removed from each genus in the phylogeny, from one species to the whole genus; 3) ED scores for the remaining species were calculated from the phylogeny; and 4) ED scores of removed species were imputed from their remaining congeners or confamiliars using each of the three methods. All four steps were repeated 100 times for each of the reference phylogenies. We ran linear regressions to test how well the ED scores from each of the three imputation methods predicted the reference ED scores.

To examine how the imputation methods performed when progressively fewer congeners and confamiliars were present in the phylogeny, we also calculated the proportion of the reference ED captured by each imputation method (i.e. imputed ED divided by reference ED) for all simulations. We then ran linear regressions of the proportion of reference ED captured by each imputation method against the proportion of that species’ genus or family retained in the phylogeny at the point of imputation. Under ideal performance of the imputation method, we would expect the slope of the regression line to be 0 (i.e. no change in proportion of reference ED predicted) and for most of the points to lie on a horizontal line at y = 1 (i.e. 1:1 proportion of imputed ED and reference ED).

During our test of imputation methods, Tonini et al. published a distribution of 10,000 species-level squamate phylogenies [35] (with 9,755 species; ~98% of species at time of its publication). In comparison to the 4,161-species (~45% complete) phylogeny of Zheng and Wiens [52], this allowed us to assess how the imputation methods perform when the phylogeny on which they are based is superseded. We used the most accurate of the three imputation methods to calculate ED values for all squamate species, based on the older phylogeny of Zheng and Wiens [52]. We then used a linear regression to test how well these imputed values predicted the median ED values from Tonini et al.’s [35] distribution of 10,000 species-level phylogenies [35]. We use the median across all 10,000 phylogenies as ED was not normally distributed.

We also compared EDGE rankings obtained from the imputed ED scores from Zheng and Wiens [52] with EDGE rankings derived from the median ED scores from the 10,000 phylogenies of Tonini et al. [35] for all species with non-Data Deficient IUCN Red List assessments and recognised in the Reptile Database [54] (3,966 species). We compared these EDGE rankings using linear regression. This allowed us to explore the stability of squamate EDGE rankings in the face of changes in available phylogenetic data.

### Identifying phylogenetic conservation priorities in the face of uncertainty

There are multiple competing phylogenies for each taxonomic group, and new phylogenies are continually being refined. We therefore need a prioritisation method that highlights a robust set of species on which to focus our efforts, where “robust” means least likely to experience a shift in ED score with future alterations to the phylogeny. We thus used the most accurate of the three imputation methods, in conjunction with published time-calibrated phylogenies for each clade, to calculate ED scores and EDGE prioritisation rankings for four groups: amphibians, birds, mammals, and reptiles. Extinct and invalid species (i.e. those not recognised by the taxonomic authorities adopted here [6,53,56,58,59]) were removed from each phylogeny prior to the calculation of ED to ensure it was not underestimated.

As reptiles are paraphyletic with the omission of birds, and no complete reptilian phylogeny exists, ED scores were calculated for the three reptilian orders from separate phylogenies. For crocodilians, ED values were calculated from Shirley et al.’s 2014 phylogeny [55]. For testudines, ED scores were calculated from Hedges et al.’s 2015 ‘megaphylogeny’ [42] cropped to the root of the testudine clade. Though two alternative phylogenies were available for testudines [43,44], one includes numerous extinct species and taxonomic discrepancies [44], while the other is not time-calibrated [43], which is essential for calculating ED scores.

For squamates, a distribution of 10,000 ED values were generated from Tonini et al.’s [35] 10,000 phylogenies, from which we took the median score for each species. We followed the taxonomy of the Reptile Database as of 15/04/2016 [54]. After calculating ED and EDGE scores separately for testudines, crocodilians and squamates we applied the ‘EDGE species’ criteria to each clade individually (species must be above median ED of their clade and be in a ‘threatened’ IUCN Red List Category (Vulnerable and above) to be considered EDGE species [12]), then combined all three groups of ‘EDGE species’ to identify the top 100 ranking EDGE species for reptiles as a whole.

For amphibians, ED values were generated from Pyron’s 2014 phylogeny [41] and imputed for all species absent from the phylogeny using the best performing imputation method. Our taxonomy followed Frost’s Amphibian Species of the World as of 01/02/2016 [56].

We calculated a distribution of 100 ED values for all mammal species, including those missing from the phylogeny, using Kuhn et al.’s [57] open access sample of 100 fully-resolved mammal phylogenies, and used the median ED value to generate EDGE scores. We did not use the recently published mammal phylogeny of Faurby & Svenning [50] to calculate ED scores as their phylogeny was constructed by prioritising topology over branch length accuracy [50], the latter being critical for ED calculation. We followed IUCN Red List taxonomy for all assessed mammal species and, for species absent from the Red List taxonomy, we referred to Wilson and Reeder [58].

We followed Collen et al. [19] by substituting imputed ED scores for published divergence time estimates for two highly distinct species comprising monotypic genera that are absent from the phylogeny (*Laonastes aenigmamus* and *Pseudoryx nghetinhensis*; see S2 dataset for scores and references). We included *P. nghetinhensis* in the calculation of Bovine ED scores to ensure we controlled for all missing species in the family. The use of the divergence times for these species, or their ‘terminal branch length’, can be considered a conservative estimate of their ‘true’ ED, as terminal branch lengths are the minimum guaranteed contribution to the ED score of any species [11,19,27].

For birds we calculated ED scores from the revised distribution of phylogenies of Jetz et al. [15], from which we imputed ED for missing species using the best performing method. We followed the taxonomy of BirdLife International’s taxonomic checklist 8.0 [59] and removed invalid species from the phylogeny before calculating ED.

We aimed to identify ‘robust’ high priority species in an EDGE framework; those which, in the absence of changes in ‘GE’ (i.e. IUCN status) or taxonomic inflation, are likely to remain high priority species irrespective of improved phylogenetic coverage. Depending on the nature of the phylogenetic data, the mechanism for identifying these ‘robust’ high priority EDGE species varied. For mammals, birds and squamates, for which ED was calculated from a distribution of relatively (>90%) complete phylogenies, we consider species which are present in the top 100 ranks across all phylogenies in the distribution as robust priority EDGE species. These are species that are invariably high-ranking when incorporating the available phylogenetic uncertainty.

However, as ED for amphibians, crocodilians and testudines was calculated from single consensus phylogenies with a large proportion of missing species (>20%), we developed separate criteria for identifying robust priority EDGE species. We assumed robust priority EDGE species to be top 100-ranked species for which all congeners or—for monospecific genera—confamiliars are present in the respective phylogeny. These are the cases with minimal uncertainty, for which no ED scores in the genera (or family) were imputed; they would only change if new species were described. The ED, and therefore EDGE, scores for these species are least likely to change with increased genetic coverage (assuming the absence of changes in ‘GE’). This assumption is supported by analyses presented in S1 Text.

## Results

### Imputation of Evolutionary Distinctiveness for missing species

For all three imputation methods (‘original’, ‘simple’ and ‘new’), imputed ED scores of species removed from phylogenies were significant predictors of the reference ED score when imputing at a genus level (all p < 0.001) and family level (all p < 0.001) for all three phylogenies (models were run separately for each taxonomic group, see Table 1). Of the three imputation methods, the imputed ED scores calculated using the new method captured substantially more of the variance in the reference ED scores (variance explained increased by an average of 59% compared to the original method, and 9% compared to the simple method; Table 1).

**Table 1.** Results from linear regression of imputed ED scores against reference ED scores for each species from the full phylogenies, using three imputation methods.

| Imputation<br>method | *Adjusted R*<sup>2</sup><br>(amphibians) | d.f. | p | *Adjusted R*<sup>2</sup><br>(mammals) | d.f. | p | *Adjusted R*<sup>2</sup><br>(reptiles) | d.f. | p |
| --- | --- | --- | --- | --- | --- | --- | --- | --- | --- |
| Original | 0.2607 | 322697 | <0.001 | 0.4332 | 479772 | <0.001 | 0.3808 | 410683 | <0.001 |
| Simple | 0.4231 | 322697 | <0.001 | 0.5589 | 479772 | <0.001 | 0.5419 | 410683 | <0.001 |
| New | 0.4941 | 322697 | <0.001 | 0.5699 | 479772 | <0.001 | 0.5899 | 410683 | <0.001 |

Regressing the proportion of the reference ED captured by the imputed ED against the proportion of each genus—or family—remaining in the phylogeny also indicated that our new imputation method is the most accurate (with most of the points centring around the y = 1 line; Fig 1), irrespective of the proportion of the phylogeny (as indicated by the zero slope; Fig 1). In this case we ran the model on data from all taxonomic groups combined, to be able to easily obtain a visual comparison of the slope and intercept of the models. Across all taxonomic groups combined, the new method overestimated reference ED scores by an average of 0.8% when imputed from congeners, and underestimated ED scores by an average of 0.2% when imputed from confamiliars. Underestimation increased when ED scores were imputed using the original method to an average of 31.3% when imputed from congeners, and 47.8% when imputed from confamiliars. The simple method overestimated ED by an average of 38% when imputed from congeners and 39.1% when imputed from confamiliars. Thus, we implemented our new method when imputing ED values for missing species in all further analyses.

**Fig 1.**
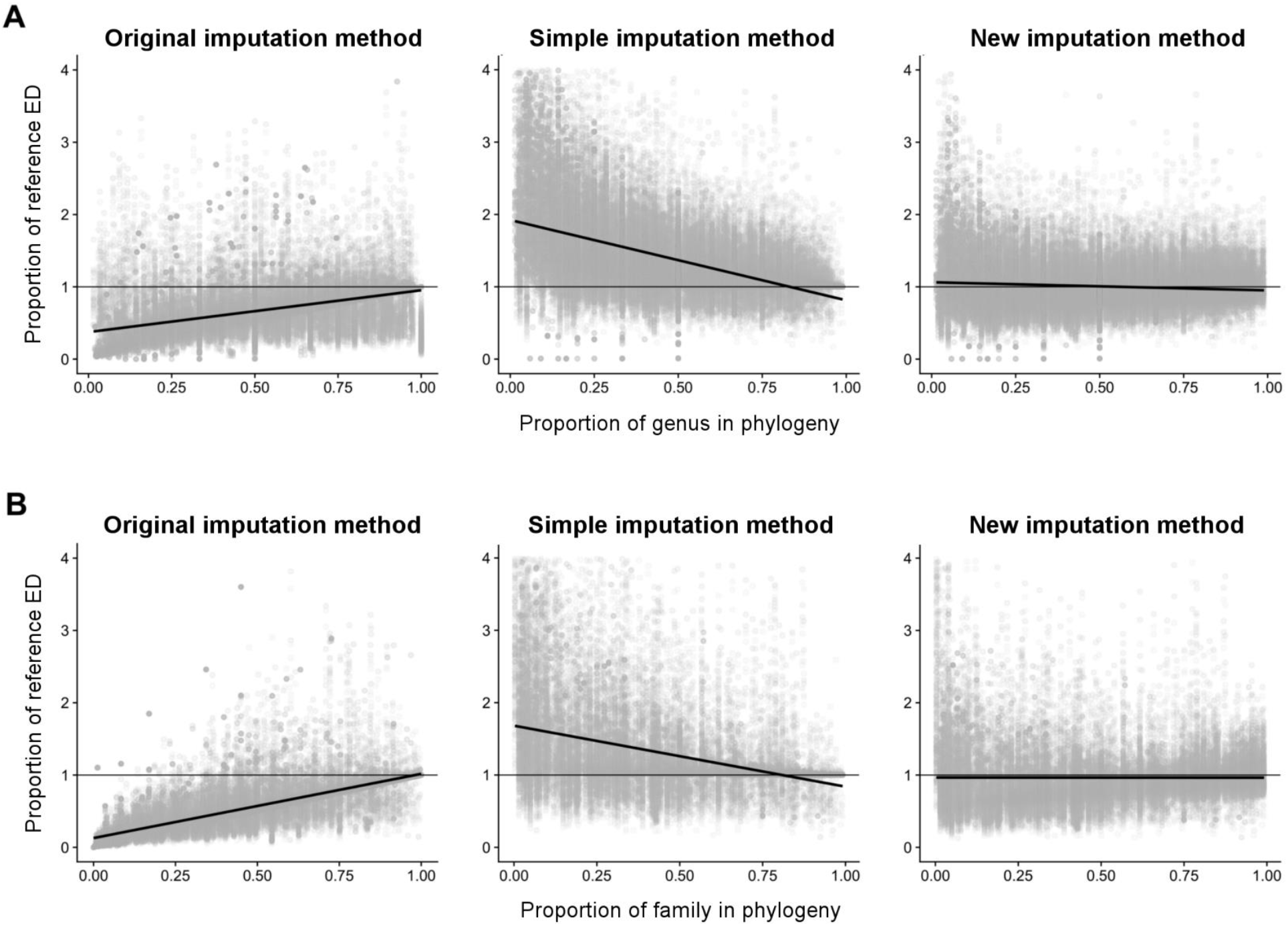
Relative performance of the three ED imputation methods. Proportion of reference ED captured when ED is imputed from: (A) congeners and (B) confamiliars for all phylogenies combined. Horizontal thin black line represents a line with an intercept of one (signifying 100% correspondence to original ED score), and a slope of zero (no change in proportion of reference ED predicted); the ideal performance of an imputation method. The thicker black line shows the modelled relationship between proportion of species removed and proportion of reference ED score estimated. Grey points represent one species from one iteration.

We then assessed the performance of imputed ED scores for squamates generated from the 4,161 species phylogeny of Zheng and Wiens [52] as predictors of the median ED scores from the newer distribution of 10,000 phylogenies of Tonini et al. [35]. We found that the imputed ED scores were a significant predictor of the median ED values generated for each species from the distribution of phylogenies (Adjusted *R^2^* = 0.5698, d.f. = 9,576, p < 0.0001; Fig 2A). EDGE rankings from imputed ED scores were also a strong predictor of EDGE rankings from the median ED of the newer distribution of 10,000 phylogenies (Adjusted *R^2^* = 0.8228, d.f. = 3,964, p < 0.0001; Fig 2B). Seventy-three of the top 100 EDGE-ranked squamate species identified using the median ED scores from the newer Tonini et al. [35] phylogeny were also in the top 100 EDGE ranking species using imputed ED scores from the phylogeny of Zheng and Wiens [52]. Of the top 50 EDGE ranking species using the median ED scores, 44 were returned in the top 100 EDGE ranks using the imputed ED scores.

**Fig 2.**
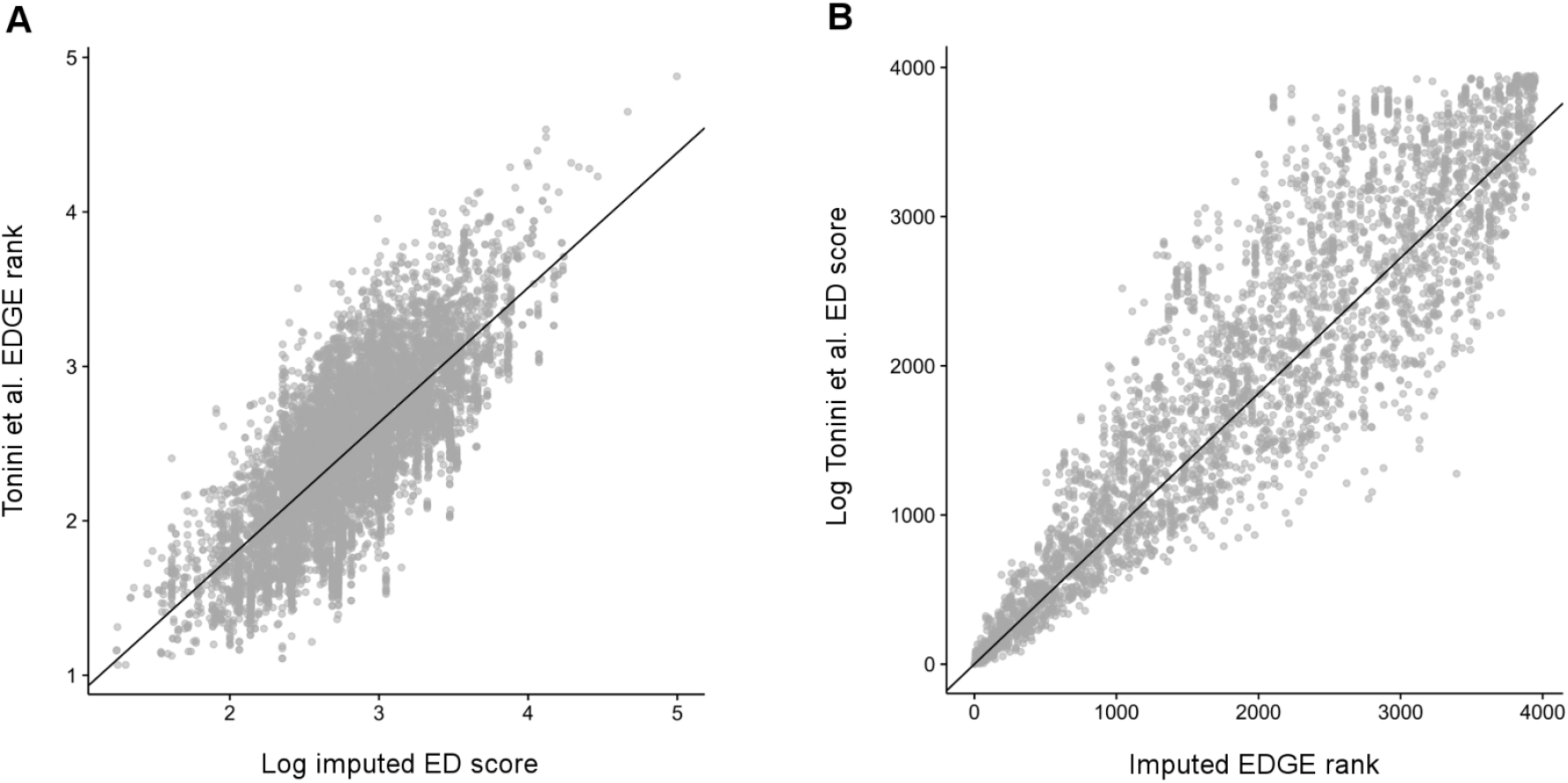
Comparison of imputed squamate ED and EDGE scores with fully phylogeny-derived ED and EDGE scores. (A) ED scores for all squamates, imputed from the 4,161-species phylogeny of Zheng and Wiens [52] using our new imputation method, against the median ED scores from Tonini et al. [35]; (B) EDGE ranks for all squamates with non-Data Deficient Red List assessments, calculated from our imputed ED scores against EDGE ranks calculated from the median ED scores from Tonini et al. [35]. Solid black line in each plot shows modelled relationship.

### Identifying robust phylogenetic conservation priorities in the face of uncertainty

We used available phylogenies and the novel imputation method to estimate ED scores for all 33,781 species of amphibians, birds, mammals and reptiles known as of February 2016 (Fig 3). When combined with available IUCN Red List assessments, we estimated the EDGE rankings for 23,387 tetrapod species (~69% of described species; Table 2 and Fig 3; all ED and EDGE scores in S2 Dataset).

**Fig 3.**
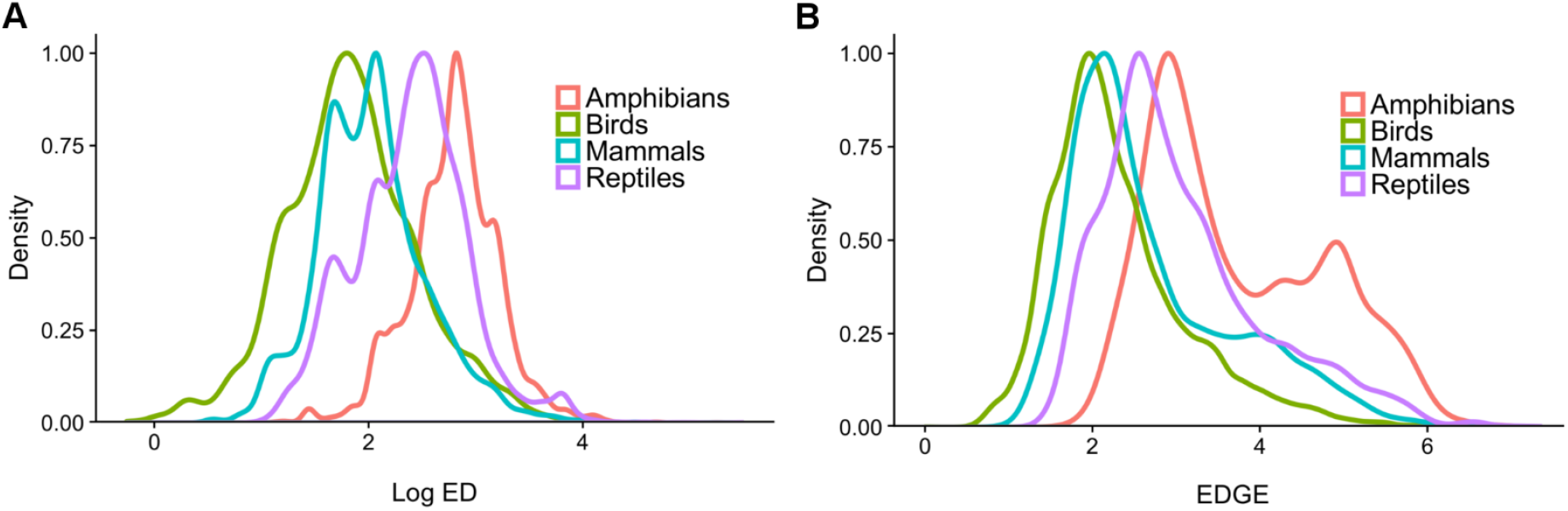
The distribution of ED and EDGE across tetrapods. Density distributions, scaled to a maximum value of 1 for (A) ED for all tetrapods using available phylogenies and imputation and (B) EDGE for all tetrapods with IUCN Red List assessments.

**Table 2.** Species richness, number of species with EDGE scores, median, maximum and total ED (in millions of years, MY) estimated for all tetrapod groups.

| Clade | No. of species<br>(as of<br>01/02/2016) | No. of<br>imputed<br>species | No. of<br>species<br>with<br>EDGE<br>scores | Median ED<br>(MY) | Total ED<br>(MY) | Maximum<br>ED (MY) |
| --- | --- | --- | --- | --- | --- | --- |
| Amphibians | 7,488 | 4,213 | 4,615 | 16.4 | 133,412 | 191.0 |
| Birds | 10,451 | 458 | 10,218 | 5.9 | 78,162 | 72.8 |
| Mammals | 5,451 | 536 | 4,619 | 7.5 | 47,943 | 89.5 |
| Reptiles | 10,391 | 502 | 4,205 | 11.3 | 131,044 | 242.9 |
| **All tetrapods** | **33,781** | **5,709** | **23,657** | **8.0** | **390,561** | **242.9** |

We estimated the first ED values for all 10,391 reptiles and the first EDGE scores for the 4,205 reptile species with non-Data Deficient IUCN Red List assessments. 9,889 species were present across the three phylogenies used to calculate ED values for reptiles, with the scores for the remaining 502 species imputed from related species present in the phylogenies.

Testudines have a higher median ED (31.0) than crocodilians (13.4) and squamates (11.1). The reptile— and tetrapod—with the highest ED is the tuatara (*Sphenodon punctatus*; median ED = 242.9); classified as Least Concern by the IUCN. 29 of the top 100 EDGE reptiles are testudines, 68 are squamates, and three are crocodilians. The highest ranked EDGE testudine, reptile, and tetrapod is *Erymnochelys madagascariensis*, a critically endangered freshwater turtle endemic to Madagascar, which we estimate to have an ED of 96.8 and an EDGE score of 7.35. The highest ranked EDGE squamate is *Xenotyphlops grandidieri*, a Critically Endangered blind snake from Madagascar, with a median ED of 67.7 and an EDGE score of 7.00. The Critically Endangered Chinese alligator (*Alligator sinensis*) is the highest ranked EDGE crocodilian, with an ED of 41.7 and an EDGE score of 6.53.

Pyron’s (2014) amphibian phylogeny included 3,275 valid amphibian species (~43% of described species), from which ED values for a further 4,213 species were imputed to give scores for 7,488 species in total. EDGE scores were then calculated for 4,615 species with non-Data Deficient IUCN Red List assessments. From our estimates, the amphibian with the highest ED is the Mexican burrowing toad (*Rhinophrynus dorsalis*; ED = 191.0). We estimate the top ranking EDGE amphibians to be the Critically Endangered Archey’s frog (*Leiopelma archeyi*) endemic to New Zealand (ED = 62.8, EDGE = 6.92) and the Chinese giant salamander (*Andrias davidianus*; ED = 61.2, EDGE = 6.90).

For mammals, the distribution of trees from Kuhn et al. 2011 [57] were modified to contain 4,915 valid and extant mammal species (90% of recognised mammal species). ED values were imputed for 536 species to give ED scores for a total of 5,451 recognised species and EDGE scores for 4,619 with non-Data Deficient Red List assessments. The aardvark (*Orycteropus afer*; median ED = 89.45) and duck-billed platypus (*Ornithorhynchus anatinus*; median ED = 89.25) have the highest mammalian ED scores. The highest ranked EDGE mammals are the Critically Endangered long-beaked echidnas of New Guinea: *Zaglossus attenboroughi* and *Zaglossus bruijnii* (ED = 46.56, EDGE = 6.63 for both species).

We imputed ED scores for 458 species of birds that have been described or reclassified since the publication of the phylogeny used by Jetz et al. [15], producing a total of 10,451 birds with ED scores. We estimate a lower median ED (5.9) than Jetz et al. (6.2; [15]) due to the inclusion of a large number of imputed species with low ED (median ED of imputed species = 5.7). The bird with the highest ED remains the oilbird (*Steatornis caripensis*; median ED = 72.8), and the highest ranked EDGE bird remains the giant ibis (*Thaumatibis gigantea*; median ED = 37.9, EDGE = 6.43).

We identified 19 of the 100 highest ranking EDGE reptiles to be robust priority species: 10 testudines, 2 crocodilians, and 7 squamates (Table 3). Only 14 of the top 100 EDGE amphibians are considered robust priority species and 15 of the top 100 EDGE bird species (Table 3). However, 78 of the top 100 EDGE mammal species were deemed robust priority species (the top 20 of these are shown in Table 3 and all 78 in S2 Dataset). All ED and EDGE scores from this analysis are available as supplementary material (S2 Dataset) and annually updated ED and EDGE scores will be available for download at www.edgeofexistence.org from 2018.

**Table 3.**
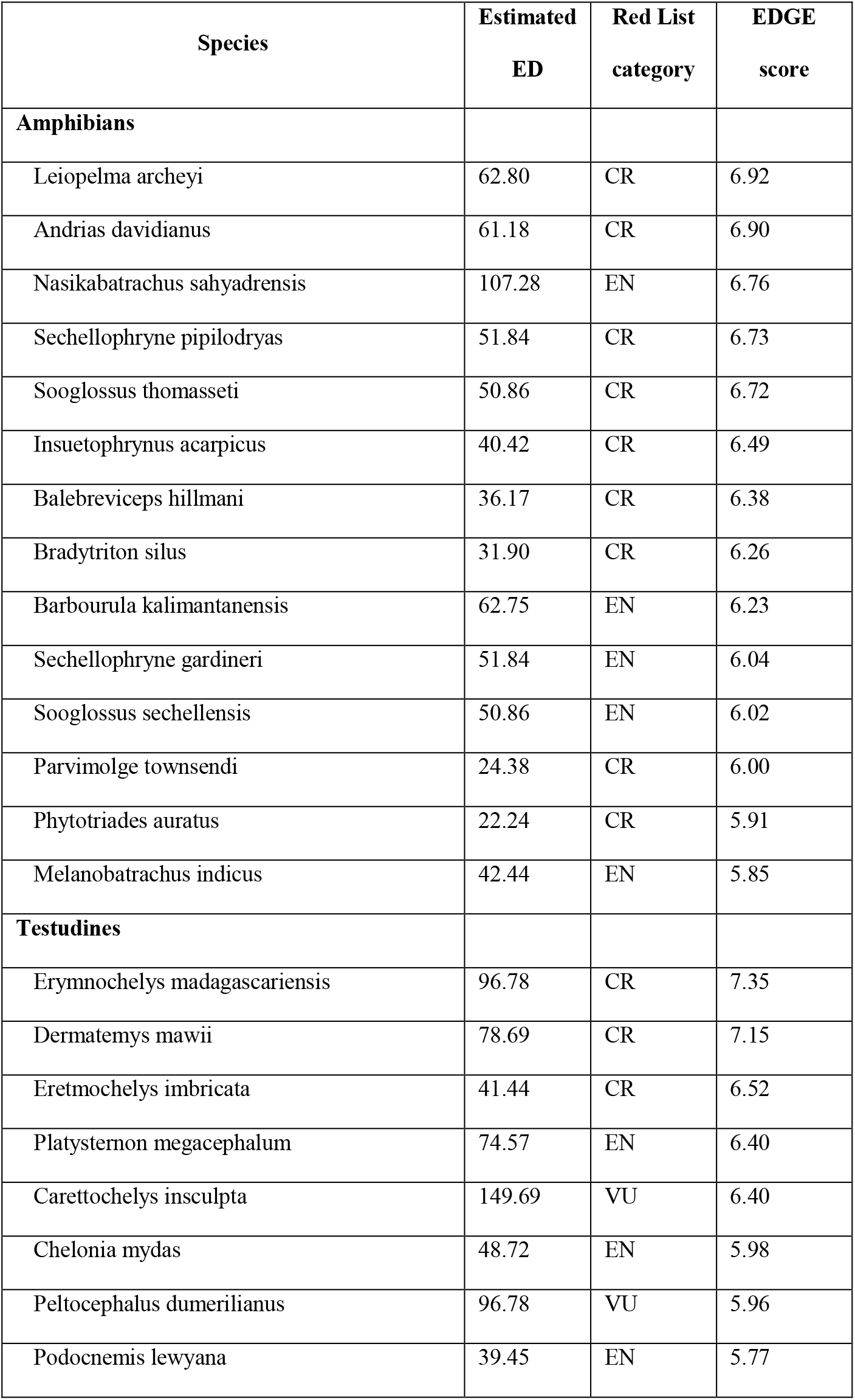
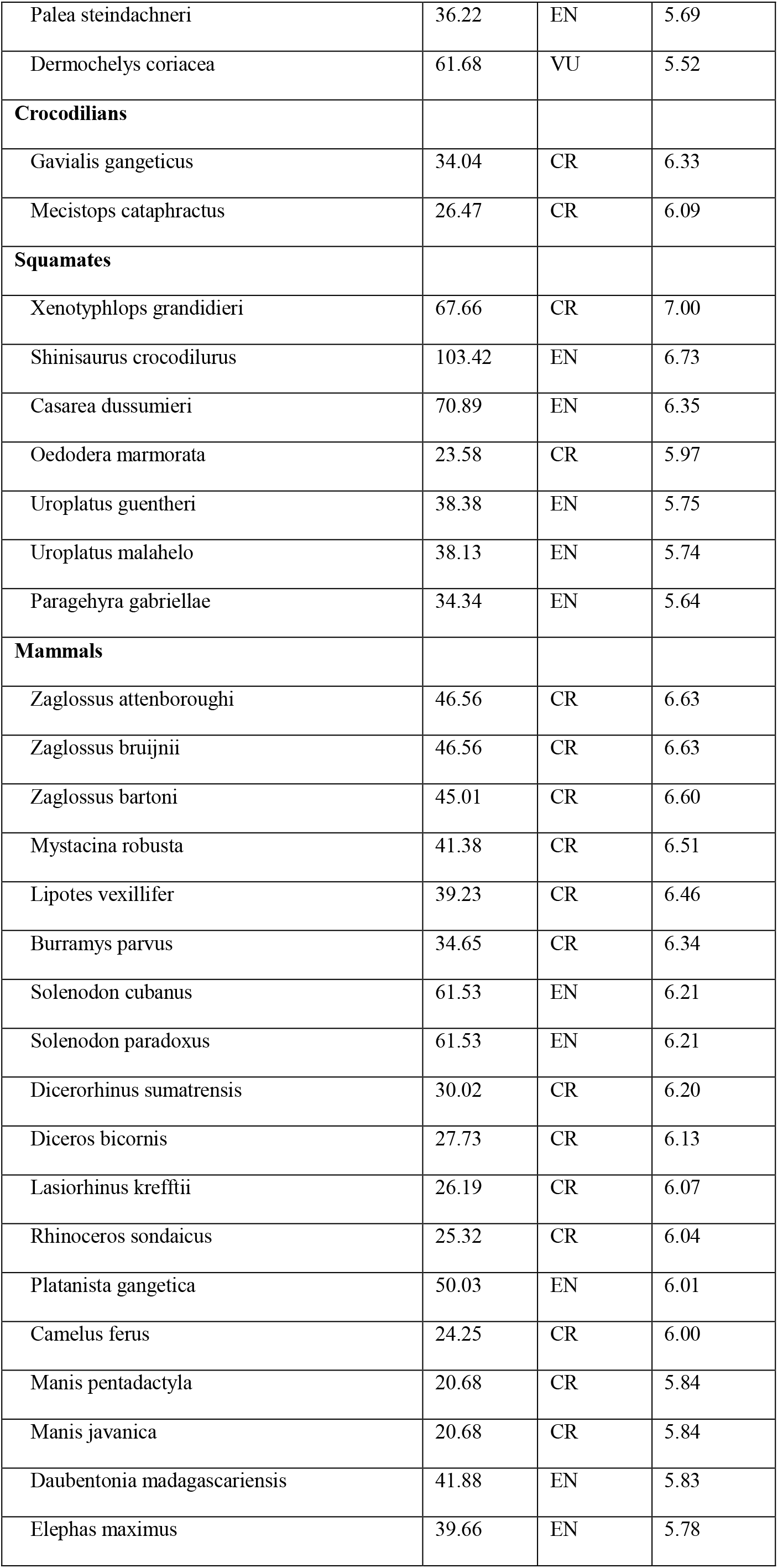
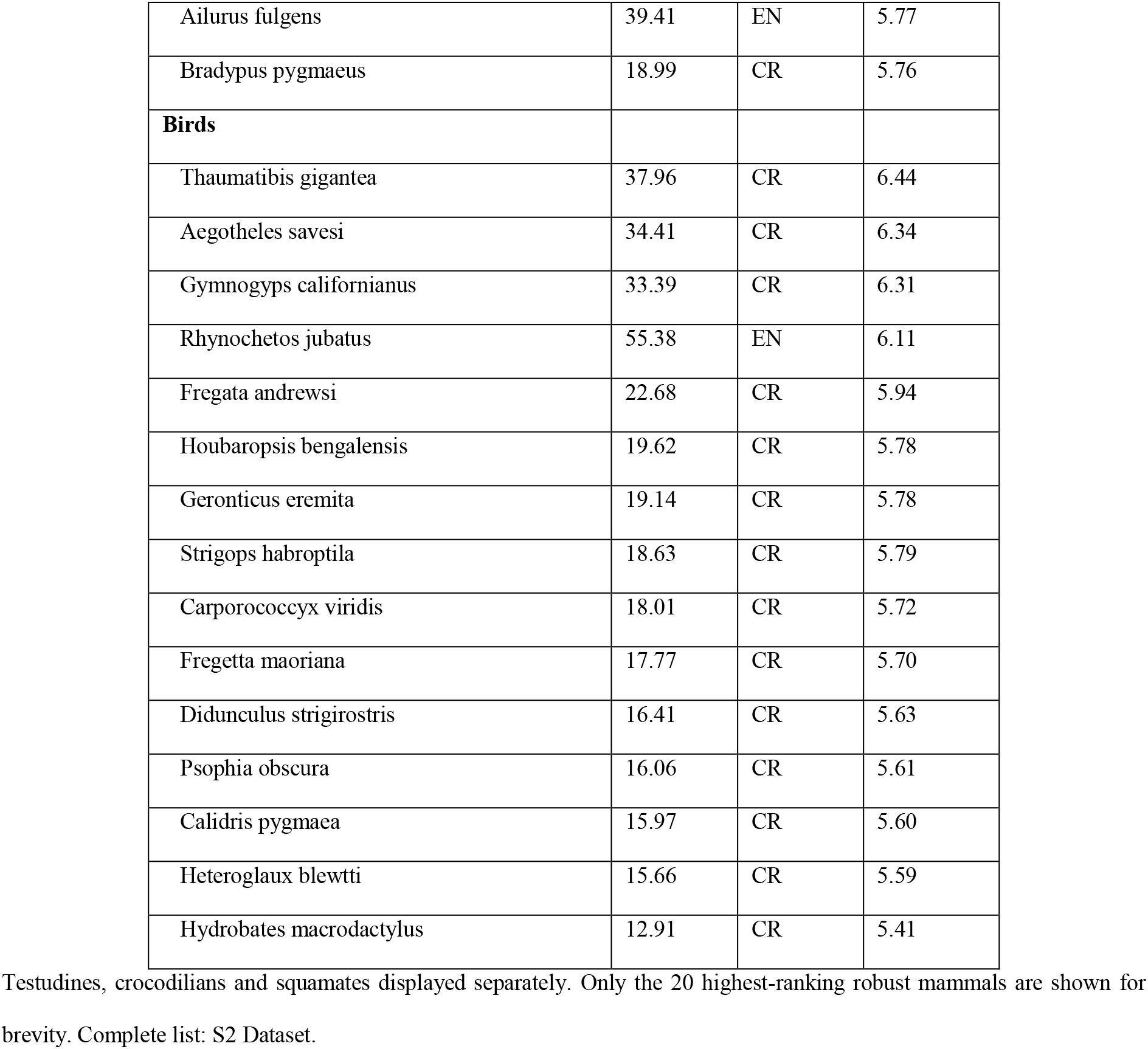
Robust top-100 EDGE species for tetrapods with IUCN Red List assessments.

## Discussion

Here, we have compared existing methods for imputing ED scores of species missing from a phylogeny with our novel method, finding that the new approach is substantially more accurate. We applied the new method to estimate ED scores for all tetrapods and, in conjunction with IUCN Red List data, updated EDGE prioritisation lists for amphibians, birds and mammals, and developed the first EDGE prioritisation for reptiles. Finally, we identified species with robust ED and EDGE scores, and thus present a practical tool for incorporating missing species to produce robust conservation prioritisations in an EDGE framework.

### Imputation of Evolutionary Distinctiveness for missing species

The rate of species descriptions means species-level phylogenies already omit new species and reclassifications by the time they reach publication. For example, the squamate phylogeny of Tonini et al. [35] contained 9,754 squamates, in accordance with the March 2015 update of the Reptile Database [54]. However, this phylogeny already omitted 446 species described between March 2015 and its publication online in April 2016 [54]. Thus, even by combining limited genetic coverage with taxonomic data [35,40,48], the rate of species descriptions—particularly in amphibians and squamates—means that the imputation of ED scores remains necessary to incorporate all species into PD-based conservation prioritisation methods.

Our novel imputation method can accurately estimate the ED of missing species for phylogenetically-informed conservation prioritisation. Though the two imputation methods previously used in analyses of ED [12,19,36] also performed well in predicting ED of missing species, the method adopted here returns values closer to the reference ED score, particularly when higher numbers of species are missing from phylogenies (Fig 1). Further, the new imputation method both increases the total ED, or PD, of the clade while reducing the individual ED scores of the closest relatives to the missing species. This more accurately reflects what happens to ED scores when new species are added to a phylogeny than in either of the earlier methods.

Further, our analysis of the squamate phylogeny shows that the imputed scores are similar to those obtained from more complete phylogenies published after the imputation was carried out. Of the top 50 EDGE squamate species obtained using the newer Tonini et al. distribution of phylogenies [35], we successfully captured 44 species in the top 100 EDGE ranks by imputing ED scores from the incomplete Zheng and Wiens phylogeny [52]. The phylogenies of Zheng and Wiens [52] and Tonini et al. [35] share much of the same genetic and fossil calibration data, thus are not independent.

Nonetheless, the strong concordance of highly ranked species when using imputed versus directly calculated ED indicates that our method can correctly identify a large proportion of the highest ranked EDGE species without needing to wait for new phylogenies to be published. As phylogenies are generally updated with incremental increases in genetic coverage (rather than new genetic data for thousands of species at a time), we anticipate that imputing from the most recent phylogeny will remain an accurate method for prioritising species for conservation action. Our results therefore demonstrate the feasibility of imputing ED scores to identify phylogenetic conservation priorities without the need for creating or updating large distributions of species-level phylogenies, which requires expertise and resources not often available to applied conservation programmes.

### Identifying phylogenetic conservation priorities in the face of uncertainty

We provide here the first estimation of ED across all tetrapods, and the first EDGE prioritisation of all tetrapods with non-Data Deficient IUCN Red List assessments. With the publication of Tonini et al.’s [35] species-level phylogeny, there is now extensive phylogenetic coverage for squamates similar to that of mammals [38,49,50], birds [15,48] and crocodilians [55] with which we underpin our analysis. However, as testudines and amphibians suffer from poor phylogenetic coverage [41,42], our scores for these clades are considered less certain. The publication of new species-level phylogenies for testudines and amphibians will provide further data against which our imputation method can be tested and refined. However, such phylogenies will likely omit newly described species for which ED must be imputed.

Our EDGE rankings for reptiles reflect the high ED and imperilment of the world’s testudines [60], with turtles and tortoises comprising 29 of the top 100 EDGE reptiles despite representing only 3.3% of reptilian species richness. EDGE rankings for reptiles are limited in coverage by the paucity of IUCN Red List assessments for the group; reptiles lag behind other tetrapod groups in extinction risk assessment, with non-Data Deficient assessments available for less than 45% of species [6,61]. Thus, as Red List assessment coverage for reptiles increases, it is likely a number of high ED species lacking assessments will enter the top 100 EDGE ranks [35].

EDGE prioritisations already existed for amphibians, birds and mammals [15,19,30], but were out-of-date in terms of both taxonomic revisions and Red List assessments. There are 26 changes to the top 100 EDGE mammal species compared to Collen et al. [19]. Species for which ED was imputed from relatives in the phylogeny comprise 21 of the top 100 ranks, 12 of which are robust priority species under our criteria. There are 19 changes to the top 100 EDGE birds since Jetz et al. [15], four of which are due to the uplisting of species on the IUCN Red List, and 15 of which are due to taxonomic revisions [15,59]. When compared to the previous EDGE amphibian list [30], only 37 of the top 100 EDGE amphibians are retained.

The first reason for this change is that our updated list used a 2014 genetic phylogeny [41] from which we estimated our ED scores for amphibians, rather than the earlier taxonomic phylogeny developed for the original EDGE amphibian analysis [30]. The difference in phylogenies produced different ED scores for amphibians compared to the original EDGE amphibian list due to the inference of differing phylogenetic relationships. The second reason is that the description in the intervening period of more than 1,400 species between the original EDGE amphibian list and the one presented here has resulted in both the reduction of ED for many species following the addition of congeners and confamilials, and resulted in the identification of priority species unknown to science at the time of the original EDGE list (e.g. *Leptolalax botsfordi*, described 2013 [62]).

Notable patterns in the priority EDGE species presented here reflect imminent threats facing certain regions or clades. For example, 47 of the top 100 EDGE amphibians are from Central and South America (compared to 55 of the top 100 in the original amphibian EDGE list [30]), reflecting the continuing severe population declines across the region [63–65]. New top 100 EDGE birds include species now threatened by trade, hunting and persecution: two vulture species (Hooded Vulture *Necrosyrtes monachus*, White-headed Vulture *Trigonoceps occipitalis*) [66–68], the Javan Green Magpie (*Cissa thalassina*) [69,70] and the Helmeted Hornbill (*Rhinoplax vigil*) [71,72].

Our updated mammal rankings capture the continued decline of Madagascan biodiversity [10,73–75], now accounting for 19 of the top 100 EDGE mammals—more than twice as many species as in Collen et al. [19]. 15 of the 19 Madagascan priority mammals are lemurs, nine of which were not previously present in the top 100. Of the nine lemur species, five entered the list as a result of uplisting on the IUCN Red List, and four were only recently described, reflecting the high description rate of threatened lemur species [6,76,77].

The number of robust priority EDGE mammals identified is much greater than any other clade, which is indicative of the comparatively much higher genetic coverage. The mammalian phylogeny differs from those used to calculate EDGE scores of birds and squamates in that it comprises only species with genetic data, thus the level of uncertainty across the distribution is significantly lower[49,51].

In contrast, the identification of relatively few robust EDGE squamates (seven of 69 in top 100; Table 3) and birds (15), reflects the broad range of ED scores calculated from the Tonini et al. [35] distribution of phylogenies. Distributions of phylogenies are created to capture phylogenetic uncertainty and compensate for the inclusion of species with no genetic data. Thus, the small numbers of robust EDGE priorities are likely a conservative estimate. Encouragingly, all seven robust EDGE priority squamate species were identified as top 100 EDGE species using our imputed ED scores from the phylogeny of Zheng and Wiens [52]. Further, the establishment of EDGE scores for reptiles through imputation has facilitated additional conservation action for two priority EDGE reptiles (the Round Island keel-scaled boa, *Casarea dussumieri* and the West African slender-snouted crocodile *Mecistops cataphractus*) [37], both of which were identified as robust priorities. This highlights the utility of developing tools to initiate conservation action using immediately available data, rather than waiting for more complete phylogenies, which is unrealistic or impossible for the majority of clades.

Under the criteria for identifying robust species when only a single, incomplete phylogeny is available, 10 testudines and two crocodiles are considered robust priority EDGE species, with their entire genus or family being present in their respective phylogeny (Table 3) [42,55]. We consider 14 of the top 100 EDGE amphibians to be robust priority species. Though this is a small proportion of the top 100, seven of the 14 are monotypic genera and the other half are from small genera (2-4 spp.). The more speciose genera and families typically have species missing from the phylogeny—thus precluding species from meeting the robust criteria.

In conclusion, we have demonstrated that our new statistical imputation method outperforms earlier approaches, and we have used this method to create updated EDGE lists for all tetrapods, including the first ever EDGE list for reptiles. As a result, we have improved our ability to keep prioritisation rankings synchronised with advances in phylogenetic knowledge and can be confident that we are focusing conservation attention on robust and high-ranking EDGE species. This methodology opens opportunities to assess and prioritise new taxonomic groups, paving the way for conservation efforts on more neglected clades, before even more unique evolutionary history is lost forever.

## Acknowledgments

Thank you to Dave Redding and James Rosindell for helpful suggestions regarding our analyses. We thank Rachel Williams and Anthony Lowney for their useful comments and discussion. We are grateful to Jonathan Baillie, founder of the EDGE of Existence Programme, for his encouragement and support. A special thank you to the EDGE Fellows working around the world, who meaningfully put our prioritisations into practice through their conservation of EDGE species.

## Supporting information

**S1 Text. Examination of robust species assumption**.

**S2 Dataset. ED and EDGE scores**. The ED and EDGE scores for all amphibians, birds, mammals and reptiles, and the robust priority species from each group.

